# *In vivo* affinity maturation of murine B cells reprogrammed to express human antibodies

**DOI:** 10.1101/2023.10.20.563154

**Authors:** Yiming Yin, Yan Guo, Yuxuan Jiang, Brian Quinlan, Haiyong Peng, Gogce Crynen, Wenhui He, Lizhou Zhang, Tianling Ou, Charles C. Bailey, Michael Farzan

## Abstract

CRISPR-edited murine B cells engineered to express human antibody variable chains proliferate, class switch, and secrete these antibodies in vaccinated mice. However, current strategies disrupt the heavy-chain locus, resulting in inefficient somatic hypermutation without functional affinity maturation. Here we show that recombined murine heavy- and kappa-variable genes can be directly and simultaneously overwritten, using Cas12a-mediated cuts at their 3’-most J segments and 5’ homology arms complementary to distal V segments. Cells edited in this way to express the HIV-1 broadly neutralizing antibodies 10-1074 or VRC26.25-y robustly hypermutated and generated potent neutralizing plasma in vaccinated recipient mice. 10-1074 variants isolated from these mice bound and neutralized HIV-1 envelope glycoprotein more efficiently than wild-type 10-1074 while maintaining or improving its already low polyreactivity and long *in vivo* half-life. We further validated this approach by generating substantially broader and more potent variants of the anti-SARS-CoV-2 antibodies ZCB11 and S309. Thus, B cells edited at their native loci affinity mature, facilitating development of broad, potent, and bioavailable antibodies and expanding the potential applications of engineered B cells.

---

Antibodies can be used to treat a wide range of diseases. They are usually safe, their Fc domains confer long half-lives in the bloodstream, and they can engage a range of immune effector functions that amplify their activities. Accordingly, many *in vitro* approaches have been developed to improve the affinity of human antibodies^1^. However, *in vivo* affinity maturation of B-cell receptors (BCRs) has key advantages over phage- or cell culture-based approaches. For example, *in vitro* selection systems require discrete selection steps, usually less than 10, often without further diversification between steps. In contrast, diversification and selection are continuous and coordinated *in vivo. In vivo* competition among BCRs is also sensitive to small affinity differences^2,3^, whereas *in vitro* selection conditions are frequently less discriminating. Finally, *in vivo* but not *in vitro* selection counter-selects against BCRs with poor bioavailability because they bind self antigens, aggregate at the cell membrane, fold or express inefficiently, are readily oxidized, or are susceptible to digestion by cellular and serum proteases^4,5^. In short, the mammalian germinal center has unique properties useful for improving antibodies intended for human use.

Although it is straightforward to generate new antibodies in animals through vaccination and isolation of high-affinity BCRs, approaches for improving human antibodies *in vivo* are more limited. Transgenic mice have been generated that express a mature or progenitor form of an HIV-1 broadly neutralizing antibody (bNAb), often to test a vaccine strategy^6–9^. Cells from these mice can be adoptively transferred to congenic host mice at physiological frequencies and expanded with appropriate immunization to improve the bNAb transgene^10,11^. However, a new mouse must be developed for each target antibody, requiring a long developmental cycle that makes it impractical as means of improving current antibody therapies. Moreover, this approach cannot be extended to non-human primates, advantageous for their longevity and similarity to humans. Finally, only a single antibody, rather than, for example, combinations of diverse antibodies or libraries of antibody variants, can be introduced in this manner.

Recently, a number of studies reported adoptive transfer of B cells engineered with CRISPR/Cas9 to express HIV-1 bNAbs^12–16^ or neutralizing antibodies targeting other human pathogenic viruses^13^ (**Extended Data Fig. 1**). A common approach for doing so introduces a single cassette expressing an exogenous light-chain variable and constant gene with a heavy-chain variable gene. This cassette is typically inserted in an intron immediately downstream of the native VDJ-recombined variable gene. When cells so edited are transferred to wild-type mice, they expand and generate neutralizing serum that reflects the potency of the original antibody. However, somatic hypermutation in these engineered cells is less efficient than that observed with unmodified cells *in vivo*^14,15,17^. Critically, useful *in vivo* affinity maturation has not been reported with these systems, implying some loss of function in these engineered cells. Fully functional B cells with programmable paratopes can be useful as scientific tools, enable more precise and responsive vaccine models, facilitate *in vivo* improvement of antibody therapeutics, and establish a foundation for adaptive B cell-based therapies.

Here we present a strategy for reprogramming primary B cells that retains their ability to affinity mature in vaccinated animals, enabling *in vivo* improvement of several antibodies. Specifically, we directly replaced the heavy- and light variable chains of murine B cells with those of human antibodies, preserving the organization and regulation of the B-cell receptor locus. This ‘native-loci’ editing approach resulted in potent neutralizing plasma and efficient, site-appropriate somatic hypermutation in response to antigen. Affinity maturation of edited cells facilitated identification of more potent and bioavailable variants of the HIV-1 broadly neutralizing antibody 10-1074. We further validated this approach by markedly improving the potency and breadth of the SARS-CoV-2 neutralizing antibodies ZCB11 and S309 against more recent Omicron variants. Thus, native-loci editing enables efficient *in vivo* maturation of human therapeutic antibodies.

## Results

### Introducing exogenous human heavy- and light-chain variable segments into their respective native loci

We sought to develop and optimize a general approach for directly replacing recombined heavy and light variable chains of mature B cells with those of human bNAbs. To do so, we first determined if homology-directed repair templates (HDRTs) with homology arms complementary to the promoter of the 5’-most heavy-chain variable (VH) segment (VH7-81 in humans), and immediately downstream of the 3’-most JH segment (JH6 in human), could template repair of a CRISPR/Cas12a-mediated double-stranded break in JH6, adapting approaches described in^18,19^ (**Extended Data Fig. 1**). We initially tested this approach with the human B-cell line Jeko-1, expressing a BCR derived from VH2-70 and JH4 gene segments. These cells were electroporated with Mb2Cas12a ribonucleoproteins (RNPs) associated with a guide RNA (gRNA) targeting JH6 and an HDRT encoding the VRC26.25 heavy-chain variable gene bounded by the aforementioned homology arms (**Extended Data Fig. 2a-c**). VRC26.25 was selected because its association with the HIV-1 envelope glycoprotein (Env) is almost wholly mediated by its heavy chain^20,21^. We observed that, using a dsDNA HDRT, 7% of cells could be edited to bind a soluble form of Env, namely a SOSIP.V7 based on the CRF250 HIV-1 isolate (CRF250-SOSIP)^22,23,24^. When adeno-associated virus 6 (AAV6) vector was used to deliver HDRTs, 38% of Jeko-1 cells could be edited. Various gRNA combinations were also evaluated, and we observed that a gRNA targeting JH6 alone afforded the highest editing efficiency (**Extended Data Fig. 2d**). An alternate strategy^15^ for introducing an exogenous promoter and heavy-chain variable region, overwriting the final JH segment, was less efficient (**Extended Data Fig. 2e**). To determine if the VRC26.25 heavy-chain fully replaced a predicted 108 kb region, from VH7-81 to JH6 of the recombined Jeko-1 chromosomal DNA, we PCR amplified genomic DNA from successfully edited cells using primers outside of the HDRT homology arms (**Extended Data Fig. 2f**). We observed an appropriately sized 5.4 kb band encoding the VRC26.25 heavy chain surrounded by parts of the VH7-81 promoter and the intron following JH6. Moreover, whole genome sequencing of VRC26.25-expressing Jeko-1 cells showed that the depth of coverage dropped by half between the VH7-81 and JH6 regions compared with unedited cells **(Extended Data Fig. 2g)**, indicating excision of this region at a single allele in most cells. Thus, native-loci editing can directly replace this large genomic region with an exogenous heavy-chain variable gene.

We then investigated whether this editing strategy could be combined with an analogous strategy targeting the kappa light-chain locus. To edit the kappa locus, we used RNPs targeting the Jκ5 regions and an HDRT with homology arms complementary to the Vκ2-40 promoter and to a region immediately downstream of Jκ5. Jeko-1 cells were electroporated simultaneously with these RNPs and those targeting the heavy-chain locus. Repair events were directed by two HDRTs encoding the heavy and light variable chains, respectively, of the bNAb 10-1074 bounded by the aforementioned homology arm (**Extended Data Fig. 2a-c**). Unlike VRC26.25, 10-1074 requires both chains to bind Env^25^. After editing with dsDNA or with AAV, 3% and 11% of cells, respectively, could bind the CRF250-SOSIP trimer, indicating that heavy and light chains were successfully introduced at their native positions (**Extended Data Fig. 2c**). Thus, a native-loci editing approach, mediated by double-stranded breaks at the 3’-most heavy- and kappa-chain J segments, can efficiently overwrite recombined human heavy- and kappa-variable genes with those of an exogenous antibody.

To determine if a similar strategy can be used to edit primary murine B cells, we compared multiple 5’ HDRT homology arms and gRNAs (**Extended Data Fig. 3a-b**). We observed efficient editing using HDRT complementary to the promoter regions of VH1-85 or VH1-64 (**Fig. 1a**). VH1-85 is the most 5’ of murine VH segments, while VH1-64 is a frequently utilized VH segment^26^. We similarly used gRNA targeting the 3’-most JH segment, JH4. Using the same principles, we targeted the kappa locus with HDRT complementary to the Vκ-135 promoter and downstream of Jκ5. These HDRTs encoded, respectively, the heavy and light chains of either 10-1074 or VRC26.25-y, a previously described breadth- and potency-improved VRC26.25 variant^19^. HDRTs were again co-electroporated as dsDNA with RNP or delivered by AAV-DJ. We observed markedly greater editing efficiencies with AAV-delivered HDRT (**Fig. 1b**), approximately 6% and 3% for VRC26.25-y and 10-1074, respectively (**Fig. 1c**). Note that some antigen-positive B cells in the VRC26.25-y-edited population may express only the VRC26.25-y heavy chain. To compare this native-loci editing approach with the intron-targeted approach described in^12–16^, we introduced a light-chain-P2A-heavy chain cassette described in^12^, into the intron following JH4 using the same JH4-targeting RNP described above. We further introduced a lesion in Cκ, limiting expression of the endogenous light chain, as in^12^ (**Fig. 1d**). Both editing efficiencies and cell viabilities after electroporation were similar to native-loci editing (**Fig. 1e-f**; **Extended Data Fig. 3c-d**). However, among successfully edited SOSIP-binding cells, native-edited cells expressed significantly more BCRs than did those edited with the intron-targeting approach, as indicated both with anti-IgM and SOSIP binding (**Fig. 1g; Extended Data Fig. 3e**). No differences in BCR expression were observed in SOSIP-negative cells that in most cases expressed the original murine BCR. We conclude that murine B cells can be edited at their native loci with efficiencies comparable to those observed with an intron-based approach, resulting in higher BCR expression.

**Fig 1:**
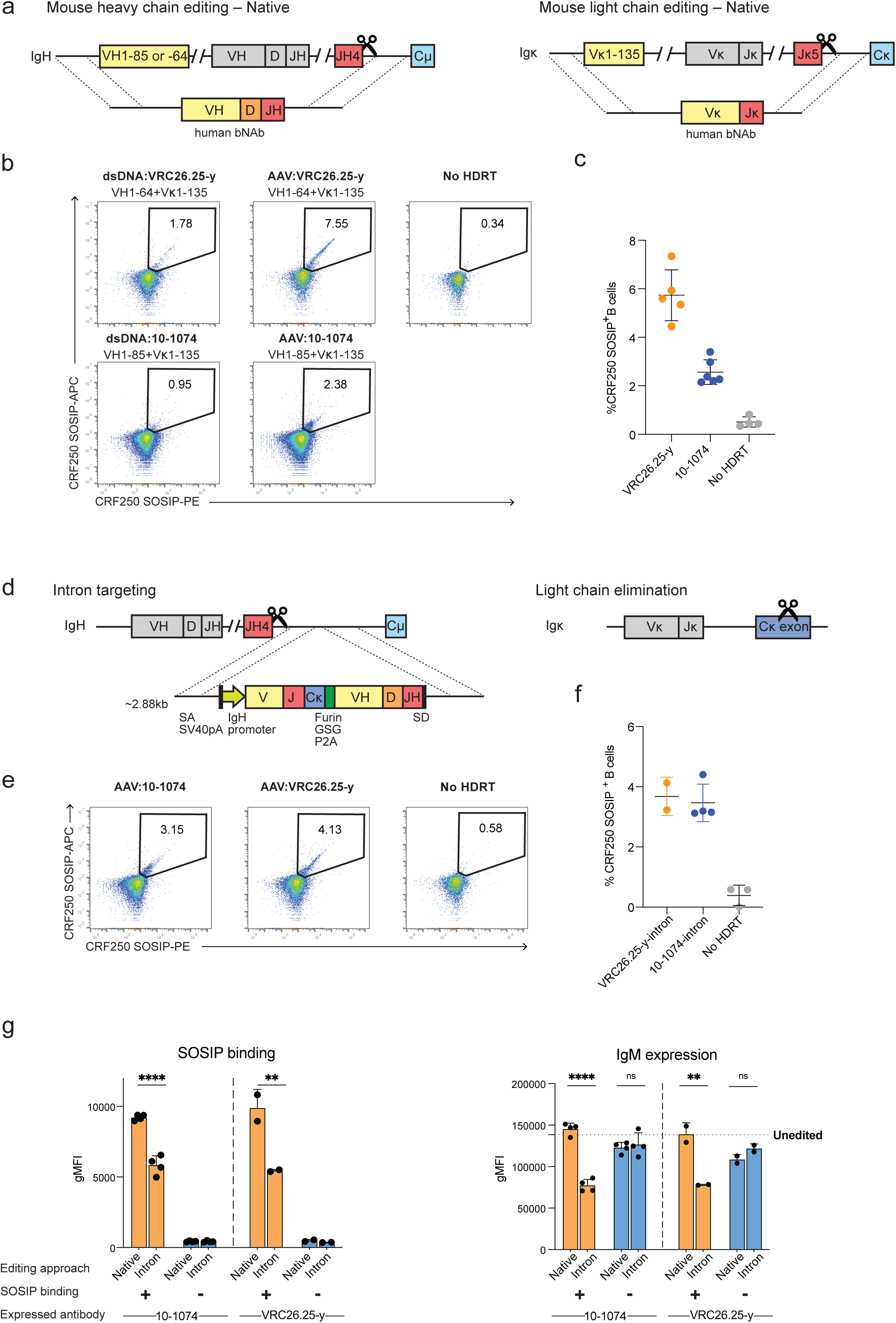
Comparison of a native-loci editing approach for engineering mouse B-cell receptors with an intron editing strategy. **a**, A representation of a native-loci editing approach in which homology-directed repair templates (HDRTs) are used to directly overwrite mouse heavy and kappa chains with exogenous heavy and light variable chains. Murine B cells were electroporated with a Mb2Cas12a ribonucleoprotein (RNP) complex targeting the murine JH4 (heavy chain, left) or Jκ5 (light chain, right) segments (scissors). HDRTs were delivered as double-stranded DNA (dsDNA) or via a recombinant adeno-associated virus (AAV-DJ). The heavy-chain HDRT (left) encodes the recombined heavy-chain variable gene of a human antibody bounded by homology arms complementary to sequence encoding the promoter and leader peptide of the VH1-85 or VH1-64 variable segments (5’ homology arm) and to the intronic region downstream of JH4 (3’ homology arm). The light-chain HDRT (right) similarly encodes a recombined kappa-chain variable region of the same antibody, with homology arms complementary to the Vκ1-135 promoter and leader-peptide sequence (5’ homology arm) and the intronic region downstream of Jκ5 (3’ arm). **b**, Primary murine B cells were edited as shown in panel (a) and analyzed by flow cytometry three days later using the same soluble HIV-1 envelope glycoprotein trimer (CRF250-SOSIP) conjugated to APC (horizontal axis) or PE (vertical axis). The bNAb introduced (10-1074 or the VRC26.25 variant VRC26.25-y^19^), the method of HDRT delivery (dsDNA or AAV-DJ), and the targets of the 5’ homology arms of the heavy (VH1-64 or VH1-85) and kappa (Vκ1-135) segments are indicated above each plot. Control cells were electroporated with the same RNP complex, without HDRT (no HDRT). Numbers indicate the percentage of cells in the SOSIP-positive gate. **c**, A summary of results from four to six independent experiments similar to those shown in panel (b), using AAV vectors to template homology-directed repair. Error bars represent SD. Note that the heavy chain of VRC26.25-y alone can bind CRF250-SOSIP. In contrast, both 10-1074 chains are necessary to bind this SOSIP. **d**, A representation of an alternative editing approach in which a bicistronic cassette is introduced in the intron between JH4 and gene segments encoding the constant domain. The cassette shown is identical to that described in^12^ and similar to that used in^13–16^. This cassette includes a splice acceptor site, an SV40 polyadenylation signal designed to terminate the murine heavy-chain VDJ message, the IgHV4-9 promoter which drives expression of an exogenous VJ-recombined human light-chain, a murine Cκ gene, a furin cleavage site, a 3-residue linker, a P2A self-cleaving sequence, an exogenous VDJ-recombined human heavy-chain gene, and a splice donor. This cassette is bounded by homology arms complementary to the intronic region, and was introduced with the same RNP used in panel (a). In parallel, the endogenous murine kappa is eliminated by NHEJ-mediated repair of a double-strand break in Cκ, as in^12^. **e**, A flow cytometry study similar to that shown in panel (b) except that the intron-targeting approach represented in panel (d) was used. **f**, A summary of results from two to four independent experiments similar to those shown in panel (e). Error bars represent SD. **g**, Three days post-electroporation, primary murine B cells edited at their native-loci as in panel (a) or at the heavy-chain intron as in panel (d) were analyzed by flow cytometry with an anti-IgM antibody and fluorescently labeled CRF250-SOSIP trimers. On the left, IgM-positive cells were gated into CRF250 SOSIP-positive (orange) and −negative (blue) populations, and the geometric mean fluorescence intensity (gMFI) from bound SOSIP trimers is presented. The corresponding IgM expression of the same cells is shown at the right. Gating strategy for these studies is presented in Extended Data Fig. 3e. Note that SOSIP-reactive cells engineered in their native loci expressed higher IgM levels than intron-edited cells, indicating higher BCR levels, whereas SOSIP-negative control cells expressed IgM comparably. Dotted line represents IgM expression observed in unedited murine B cells cultured in the same manner. Error bars indicate SD from two (VRC26.25-y) or four (10-1074) independent experiments. Significant results are indicated from left to right: ****p < 0.0001; **p = 0.0089; ****p < 0.0001; ^ns^p = 0.9389; **p = 0.0057; ^ns^p = 0.4428; two-way ANOVA with Tukey’s comparisons.

### Native loci-edited cells generate more potent neutralizing plasma after immunization

We next sought to determine whether differences in BCR expression or regulation would impact the development and maturation of B cells edited with native-loci or intron-targeting methods. B cells isolated from spleens of CD45.1 donor mice were edited with each method and adoptively transferred the same number of successfully edited B cells to CD45.2 recipient mice (**Fig. 2a**). Mice were immunized one day after transfer and at weeks 3, 5, 9 and 12 thereafter, and blood was collected between immunizations. Mice receiving B cells engineered to express 10-1074 were immunized with a 60-mer I3-01 scaffold presenting the BG505-T332N gp120 subunit of Env^27,28^. Mice receiving VRC26.25-y edited B cells were immunized with CRF250-SOSIP (**Extended Data Fig. 4a**). Plasma samples were characterized for their ability to neutralize CRF250 pseudoviruses. Plasma from mice that did not receive engineered B cells but which were immunized on the same schedule did not neutralize CRF250 pseudoviruses, indicating that responses in engrafted mice derived from the engineered cells (**Fig. 2b-c**, gray lines). The potency of response was greater with native-loci editing than with intron editing for both antibodies, reaching significance after the first (10-1074, VRC26.25-y) and third (VRC26.25-y) immunizations (**Fig. 2b-d**). These differences also extended to neutralization against HIV-1 isolates from divergent clades, significantly for the clade A isolate BG505-T332N (**Fig. 2e-f**). Notably, VRC26.25-y plasma neutralized the VRC26.25-resistant WITO isolate, consistent with the greater breadth of the VRC26.25-y variant^19^. We also monitored the viability and proliferation of successfully edited cells *in vivo*. The percent of SOSIP-positive, CD45.1+ donor cells steadily and similarly increased after each of five vaccinations in both intron-edited and native loci-edited groups of mice, rising from 5% to 40% in most cases (**Extended Data Fig. 4b**). However, compared with intron-edited cells, we observed a significantly higher percentage of SOSIP-positive donor cells among germinal center B cells in mice receiving native-loci edited cells. (**Extended Data Fig. 4c-d**). Consistent with this observation, a greater proportion of native-loci edited cells class switched to IgG (**Extended Data Fig. 4e**). We conclude that native-loci edited B cells proliferated *in vivo*, accessed germinal centers, class switched, and generated potent neutralizing plasma.

**Fig. 2:**
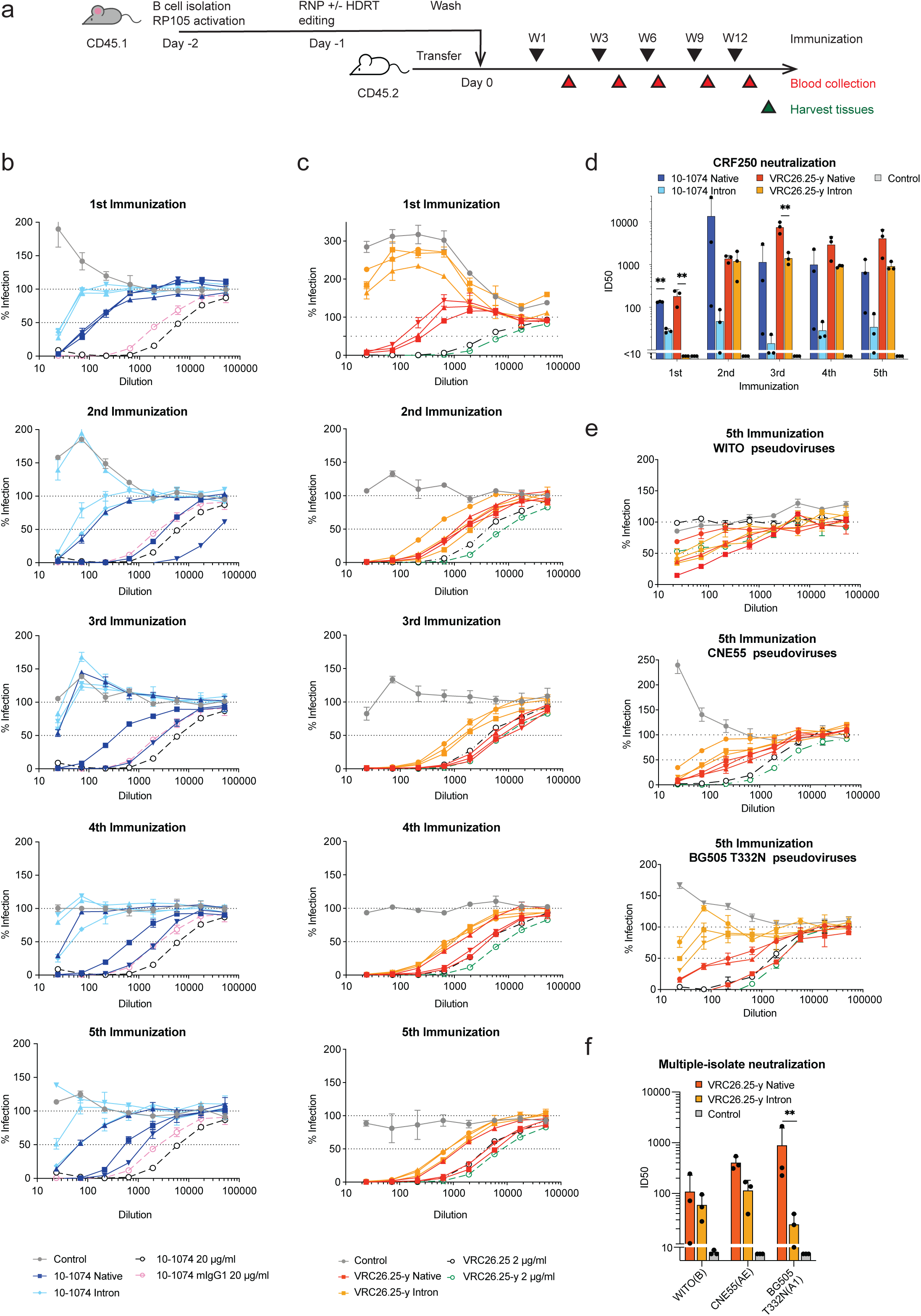
B-cells engineered with native-loci editing generate potent HIV-1 neutralizing plasma *in vivo*. **a,** The schedule of immunization and blood collection used to generate plasma and cells analyzed in panels (b-f). Murine B cells were obtained from C57BL/6 mice marked with the CD45.1 allotype, engineered with either native-loci or intron editing approaches, and adoptively transferred to CD45.2-positive recipient mice. These recipient mice were immunized with CRF250 SOSIP (VRC26.25-y) or the I3-01 BG505-T332N 60-mer (10-1074) at weeks 1 3, 6, 9, and 12 after transfer. Blood was collected at weeks 2, 4, 6, 10, and 13. **b,** Neutralization studies against the CRF250 HIV-1 isolate, homologous to the CRF250-SOSIP antigen, are shown. Panels characterize plasma after each of five vaccinations from mice engrafted with the same numbers of SOSIP-reactive 10-1074 native-loci- and intron-edited B cells. Each panel compares plasma from three native-loci-edited (dark blue) mice with three intron-edited (light blue) mice. Naïve murine plasma mixed with 20 µg/ml of 10-1074 with a murine (purple dotted line) or human (black dotted line) constant domain serve as positive controls. Negative control (gray) indicates identically vaccinated mice without adoptive transfer. Error bars indicate SEM of duplicates. **c,** A series of studies similar to those in panel (b) except native-loci introduced VRC26.25-y (orange) is compared to its intron-edited counterpart (yellow). Black and green dotted lines indicate plasma obtained from wild-type mice mixed with 2 µg/ml of wild-type VRC26.25 or the VRC26.25-y variant, respectively. Gray indicates identically vaccinated control mice without adoptive transfer. Error bars indicate SEM of duplicates. **d,** A summary of 50% inhibitory dilution (ID_50_) values from panels (b and c). Comparisons between native-loci- and intron-editing are significant after the first immunization for 10-1074 (**p = 0.0023) and VRC26.25-y (**p = 0.0049) and after the third immunization for VRC26.25-y (**p = 0.0967), as determined by mixed effects model of repeated measures followed by Tukey’s multiple comparisons. Pooled sera from mice before immunization did not neutralize CRF250 (ID_50_ < 10). Error bars represent SD for each group of three mice. **e,** A neutralization study similar to that in panel (c) except that neutralization of heterologous isolates CNE55, WITO, and BG505-T332N was measured. Error bars indicate SEM of duplicates. **f,** A summary of ID_50_ values from panel (e) indicating that each plasma retained the breadth of VRC26.25-y. Error bars represent SD for each group of three mice. Significance was determined by two-way ANOVA followed by Tukey’s multiple comparisons: **p = 0.0038.

### Efficient somatic hypermutation of native-loci edited B cells

One week after the final immunization, donor CD45.1 B cells were harvested from the lymph nodes and spleens and analyzed by next-generation sequencing (NGS) to determine the frequency of somatic hypermutation (SHM) in successfully edited cells. A higher percentage of 10-1074 or VRC26.25-y heavy- and light-chains bore coding substitutions, with more substitutions per sequence, in native-loci edited cells than in intron-edited cells (**Fig. 3a**). The frequency of amino-acid changes at each site is shown for heavy and light chains of both antibodies (**Fig. 3b**). Note that the pattern of hypermutation is different in the codon-optimized 10-1074 than it is from the donor-derived VRC26.25-y sequence, perhaps reflecting a different distribution of mutational hotspots. Alternatively, different selection pressures on the two bNAbs could impact these hypermutation patterns. The frequency of heavy- and light-chain coding substitutions was significantly greater in native-edited cells than in intron-edited cells (**Fig. 3c; Extended Data Fig. 5a**). Moreover, only native-loci edited cells exhibited a significant bias toward substitutions in the CDR region (**Fig. 3d; Extended Data Fig. 5b-c**), consistent with the pattern of naturally occurring somatic hypermutation^29^. Thus, compared with intron-targeted editing, native-loci editing resulted in significantly greater SHM that was more focused on CDR regions.

**Fig. 3:**
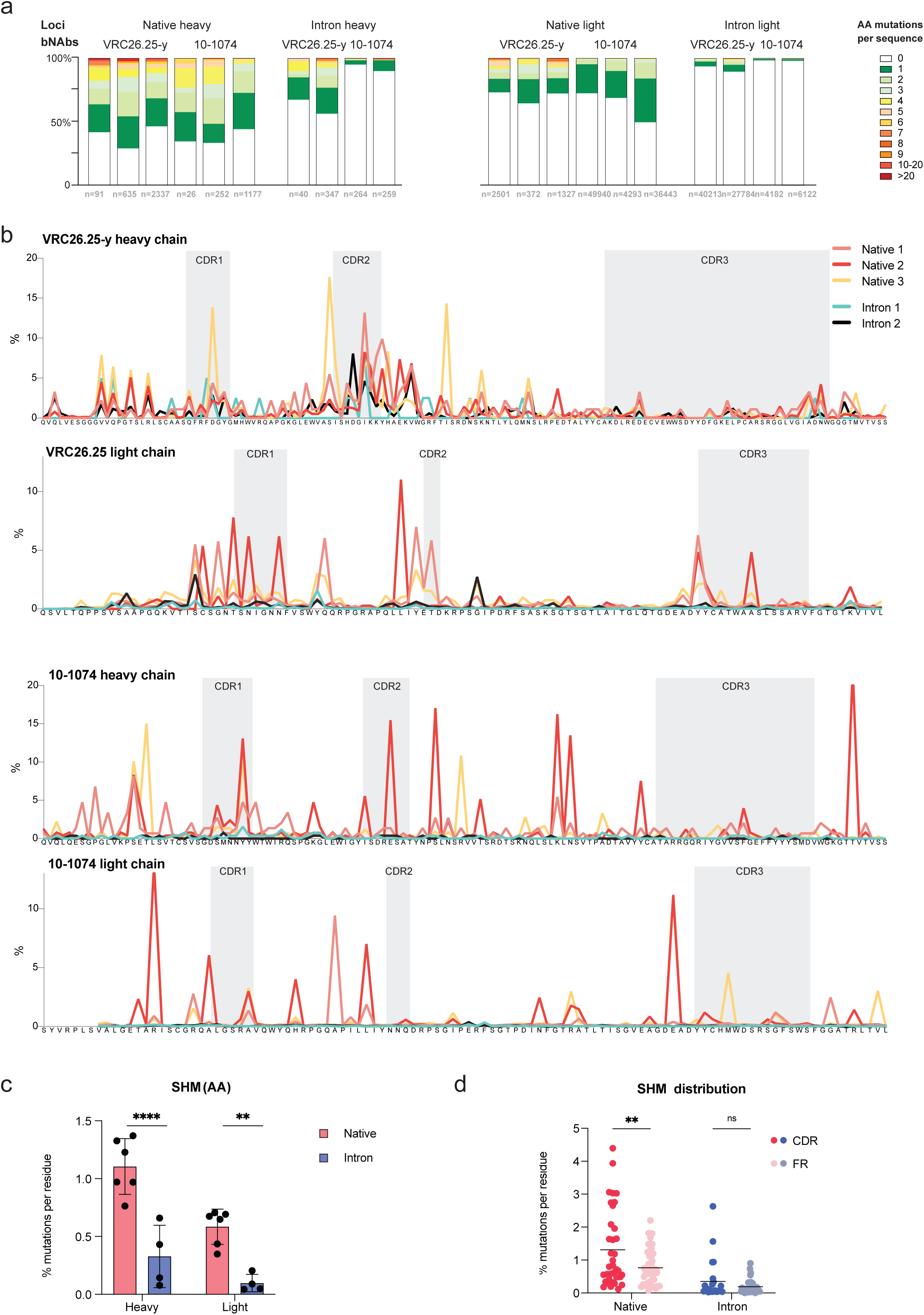
Native-loci editing enables more efficient somatic hypermutation. CD45.1 donor B cells from spleen and lymph nodes in VRC26.25-y and 10-1074 recipient mice characterized in Fig. 2 were sequenced one week after their final vaccination. **a,** The number of amino-acid mutations per sequence is represented with proportions in each library. Each bNAb and locus are indicated above the figure, and the number of unique sequences is shown below. **b,** Figure shows the frequency of amino-acid substitutions at each VRC26.25-y and 10-1074 residue, observed in three mice engrafted with native-loci-edited cells and two mice engrafted with intron-edited cells. Gray indicates heavy- and light-chain CDRs. Note the higher frequency of SHM, especially in the CDR regions, in native loci-edited mice. **c,** The percent of heavy- and light-chain amino-acid mutations is shown for native-loci (n = 6) and intron-targeting (n = 4) approaches for both antibodies. Error bars indicate SD. ****p < 0.0001; **p = 0.0033; Two-way ANOVA with Šídák’s multiple comparisons. **d,** The percent of amino-acid mutations in CDRs of both antibodies is compared to those in framework (FR) regions from native-loci and intron-edited B cells. Values for individual FR and CDR regions are presented in Extended Data Fig. 5c. Significance is indicated from left to right: **p = 0.0015; ^ns^p = 0.6465; Two-way ANOVA with Šídák’s multiple comparisons.

### VRC26.25-y- and 10-1074-expressing B cells infused into the same mouse combine to provide broader neutralization

We investigated whether native loci-edited B cells engineered to express two bNAbs could be combined to improve the breadth of the resulting plasma. A 50:50 mixture of 10-1074- and VRC26.25-y-edited cells was adoptively transferred to recipient mice and these mice were vaccinated with the CRF250-SOSIP antigen. In parallel, mice were infused with the same total number of edited cells expressing either 10-1074 or VRC26.25-y alone and similarly vaccinated (**Fig. 4a**). Plasma isolated from mice receiving both sets of edited cells neutralized viruses resistant to 10-1074 (X1632, CNE55) or VRC26-25 (JR-FL), indicating that both antibodies were present (**Fig. 4b**). In contrast, plasma from mice infused with B cells expressing a single antibody failed to neutralize viruses resistant to that antibody. We further characterized SHM in mice receiving both sets of edited cells and observed extensive hypermutation of each antibody (**Fig. 4c**). The pattern of high-frequency mutations was similar to those found in mice infused with each antibody alone, shown in Fig. 3b. We conclude that B cells engineered to express two divergent bNAbs can respond in parallel to a common SOSIP antigen, increasing the breadth of response to HIV-1 Env.

**Fig. 4:**
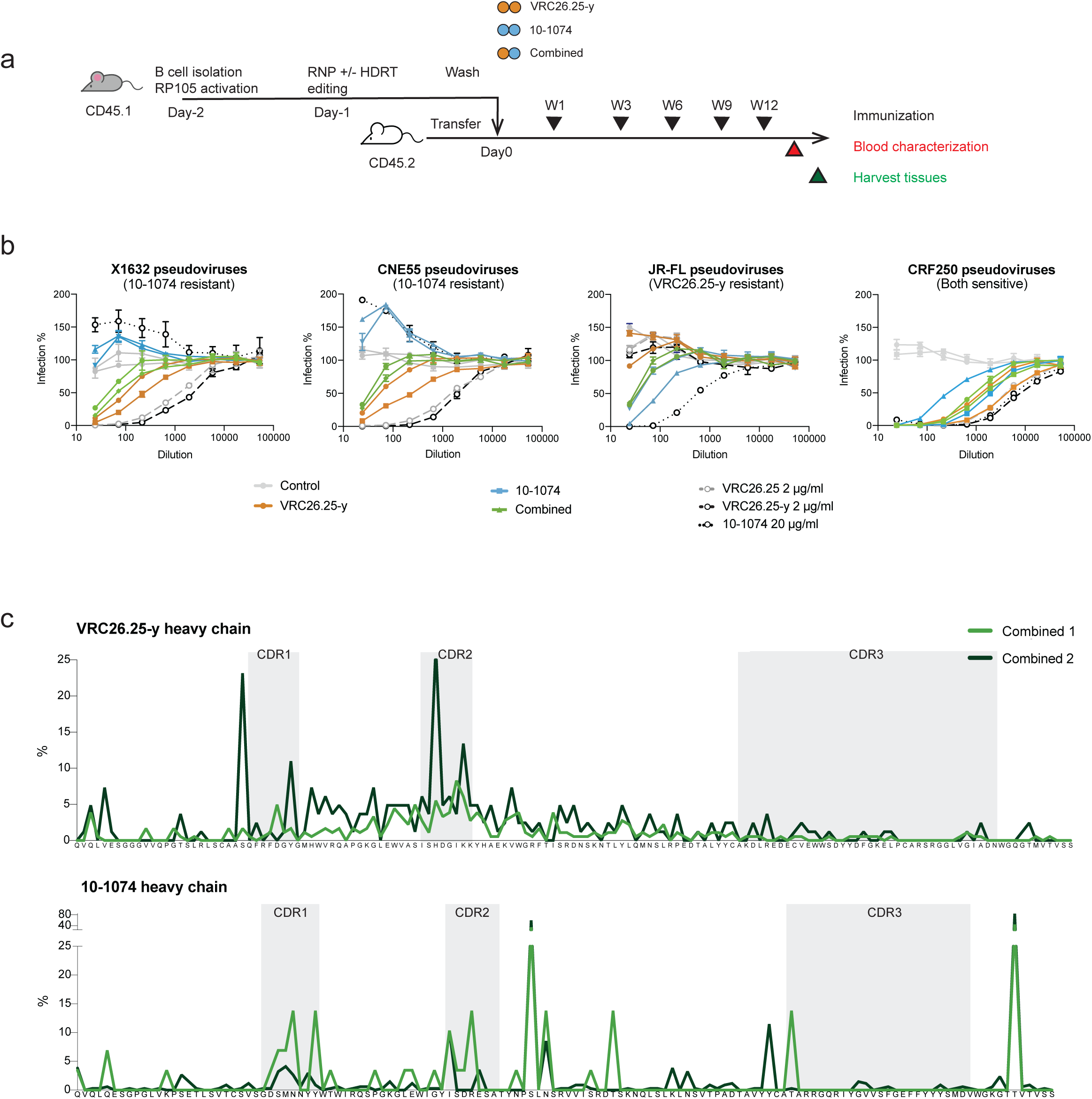
VRC26.25-y and 10-1074-expressing B cells can be combined to provide broader neutralization. **a,** The schedule of immunization and blood collection in panels (b-c). Murine B cells were obtained from C57BL/6 mice marked with the CD45.1 allotype, engineered with native-loci editing approach to express VRC26.25-y or 10-1074, and engineered cells were adoptively transferred into CD45.2-positive recipient mice, separately or in combination. Each mouse received the same number of successfully engineered cells. Mice were immunized with CRF250 SOSIPs at weeks 1 3, 6, 9, and 12 after transfer. Blood was collected at week 13. **b,** Neutralization studies against the indicated HIV-1 isolates using plasma harvested after the fifth immunization, according to the schedule in panel (a). X1632 and CNE55 are both resistant to 10-1074, whereas JR-FL is resistant to VRC26.25-y and CRF250 is sensitive to both antibodies. Each panel compares plasma from two VRC26.25-y mice (orange), two 10-1074 mice (blue), and two mice receiving both antibodies (green). Naïve murine plasma mixed with 20 µg/ml of 10-1074 with human (black dotted line) constant domain, 2 µg/ml of wild-type VRC26.25 (gray dashed line) or the VRC26.25-y variant (black dashed line), serve as positive controls. Negative control (gray) indicates identically vaccinated mice without adoptive transfer. No neutralization was observed in pooled sera from mice before immunization. Error bars indicate SEM of two replicates. **c,** The average frequency of amino-acid substitutions at each position of the VRC26.25-y and 10-1074 heavy chains in mice engrafted to express both antibodies.

### Affinity maturation of 10-1074 in wild-type mice

Phylogenetic analysis of B cells isolated from the spleens and lymph nodes of mice engrafted to express 10-1074 highlights the greater diversity of heavy-chain sequences from native-loci edited B cells compared with intron-edited cells (**Fig. 5a**). To determine if efficient somatic hypermutation in native-edited cells resulted in affinity maturation, fifteen heavy-chain sequences were randomly selected for further characterization (selected variants indicated in pink and red). Surface plasmon resonance (SPR) measurements indicate that all 15 naturally emerging 10-1074 variants bound BG505-T332N SOSIP protein with higher affinity than unmodified 10-1074, in four cases by greater than 10-fold (**Fig. 5b; Extended Data Fig. 6a**). Similarly, all of these antibodies bound surface-expressed Env (BG505-T332N-ΔCT) and neutralized BG505-T332N pseudoviruses significantly more efficiently than wild-type 10-1074. Results from all three assays represented in Fig. 5b were moderately but significantly correlated (**Extended Data Fig. 6b**), and in particular the four highest affinity variants determined by SPR were also among the most efficient in the other two assays. Notably, three of these four variants bore an S100fA mutation present in their CDR3 regions (**Fig. 5c**), suggesting that this mutation contributes to the increased affinity of these variants. In addition, we characterized all seven heavy-chain variants identified in mice receiving intron-edited cells. Four of these did not express, perhaps explained by the loss of a key disulfide bond or potentially disruptive framework substitutions. Among the remaining three, two of these 10-1074 variants did not detectably bind BG505-T332N SOSIP or surface-expressed Env, and a third variant bound and neutralized BG505-T332N pseudoviruses similarly to unmodified 10-1074 (**Fig. 5b; Extended Data Fig. 6a**). We thus observed no significant improvement of affinity in intron-edited cells, and therefore we focused our subsequent efforts on native-edited cells alone. Collectively these data show that most naturally emerging heavy-chain variants isolated from native loci-edited cells bound and neutralized more efficiently than 10-1074 itself, and that these engineered B cells affinity matured in engrafted wild-type mice.

**Fig. 5:**
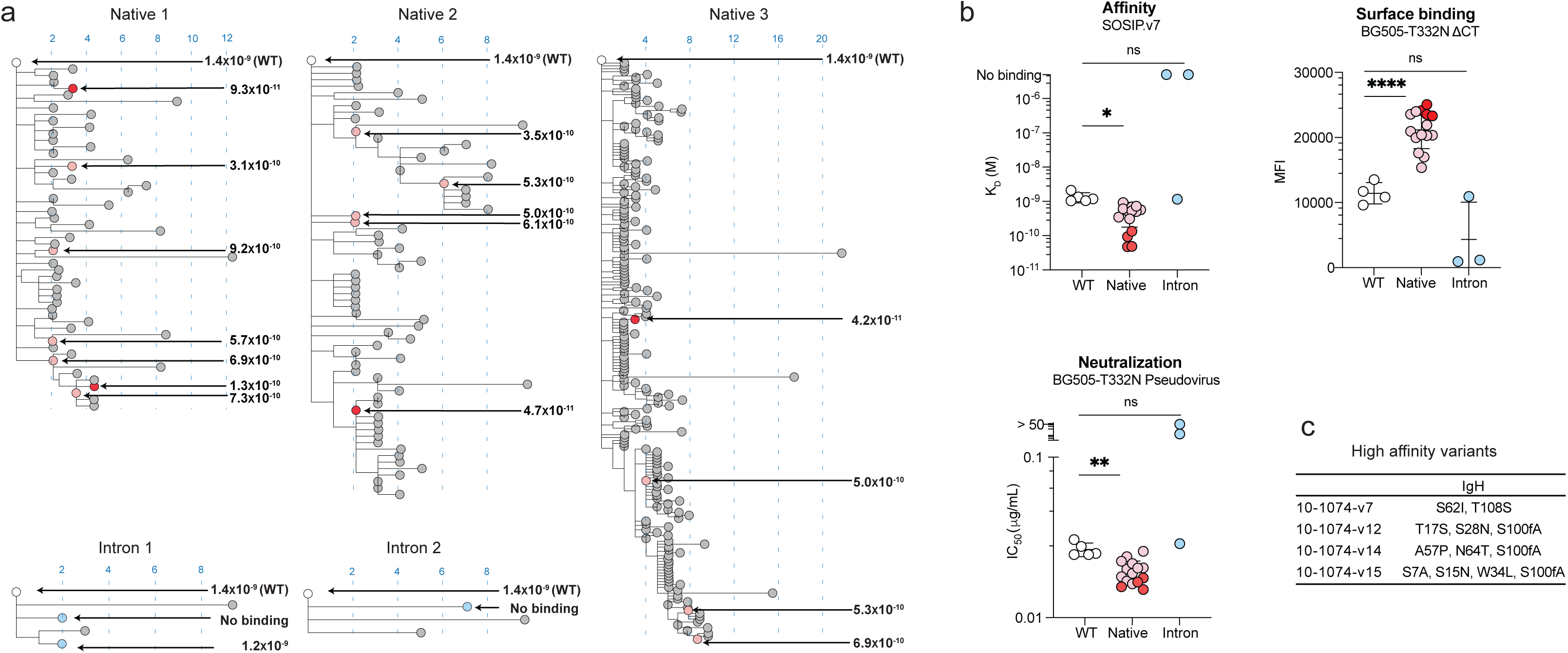
Affinity maturation of 10-1074 in wild-type mice. **a,** Phylogenetic analyses are presented of heavy-chain 10-1074 variants with multiple amino-acid substitutions isolated one week after the final immunization from mice characterized in Figs. 2 and 3. Distance from wild-type 10-1074 indicates the number of amino-acid substitutions. The affinity of unmutated 10-1074 (white) and 15 randomly selected variants (pink and red), is provided for native loci-edited cells. Red indicates four variants with affinities 10-fold higher than wild-type 10-1074. Grey indicates uncharacterized variants. Among intron-edited cells, blue indicates three characterized variants with affinity provided, while grey indicates four uncharacterized variants that did not express. **b,** Panel represents the affinity of 15 sampled native loci-edited and 3 intron-edited variants, as determined by SPR, their association with cell-expressed Env as determined by flow cytometry, and their neutralization potency. Higher affinity variants are indicated in red, corresponding to those in panel (a). Significance was determined by Welch’s ANOVA with Dunnett’s multiple comparisons: *p = 0.0159, ^ns^p = 0.2882 (affinity); ****p < 0.0001, ^ns^p = 0.2722 (surface binding); **p = 0.0018, ^ns^p = 0.5989 (neutralization). **c,** Mutations of the four high-affinity variants are listed.

### Development of more potent 10-1074 variants

The presence of S100fA in high-affinity 10-1074 variants isolated independently from separate mice suggested that mutations emerging in multiple mice would enhance the neutralization potency of 10-1074. We accordingly comprehensively characterized high-frequency mutations that emerged in more than one native loci-edited mouse (**Fig. 6a; Extended Data Fig. 7**). 32 heavy-chain mutations, including S100fA, and 16 light-chain mutations were selected based on these criteria and characterized individually for their ability to neutralize the CRF250 and BG505-T332N isolates (**Fig. 6a**). The majority of heavy- and light-chain mutations improved neutralization against at least one isolate (**Fig. 6b**). None of the mutations characterized dramatically impaired neutralization, consistent with efficient counter-selection against poorly binding variants *in vivo*. Interestingly, three of these mutations (N31D, S100fA, and Y100nF) were also found in the antibody PGT121, a V3-glycan antibody similar to 10-1074 that emerged from the same progenitor B cell of the same donor^25^. Half (16/32) of heavy-chain mutations and 31% (5/16) of light-chain mutations improved neutralization against both isolates (**Fig. 6b**).

**Fig. 6:**
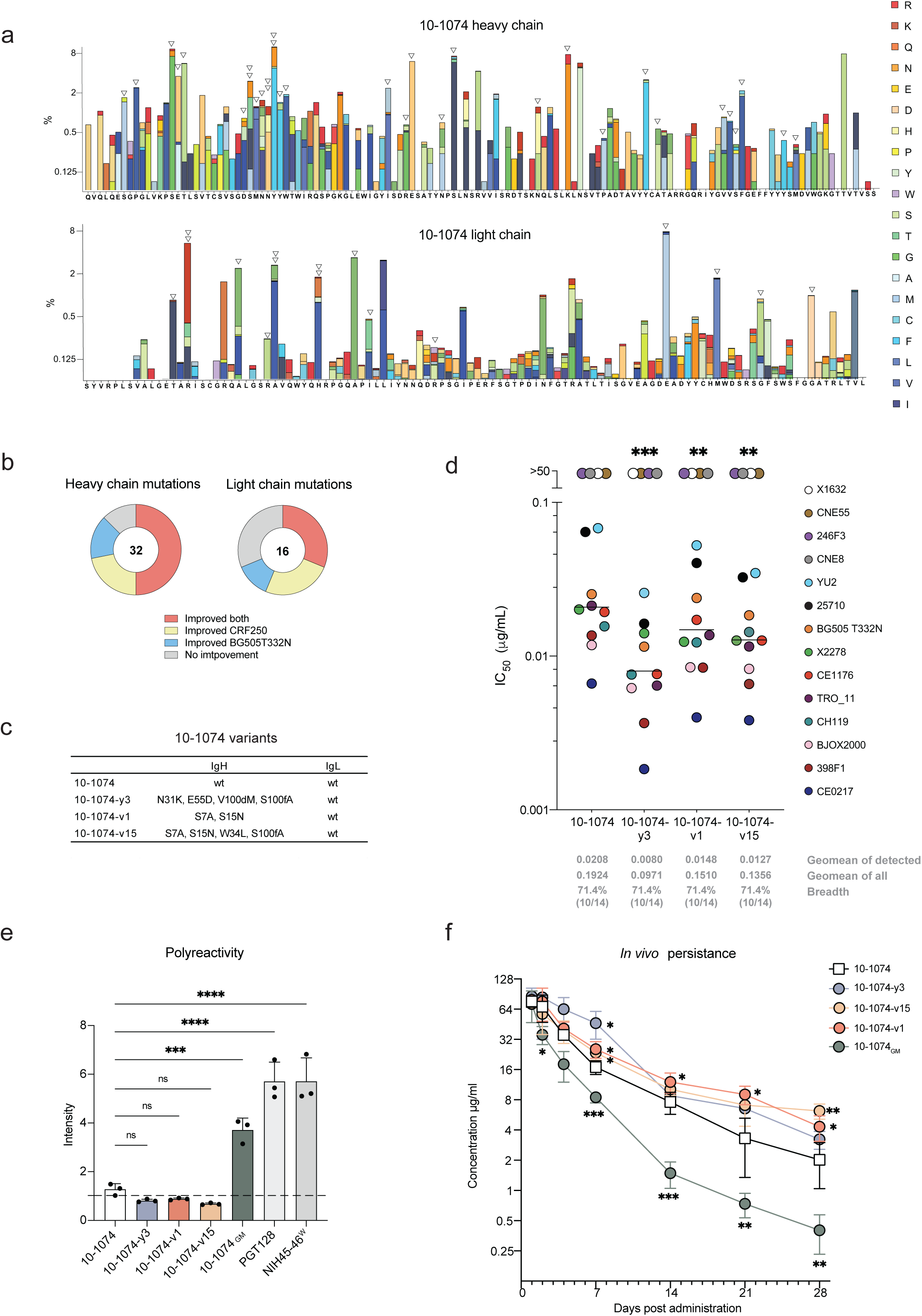
More potent and bioavailable 10-1074 variants. **a,** The frequency of heavy- and light-chain amino-acid mutations averaged from three mice engrafted with native-loci edited 10-1074 cells. White arrowheads indicate positions where the most frequent mutation emerged in more than one mouse, the criteria for further characterization. Two arrowheads indicate that the two different substitutions were present at the same position in more than one mouse, and both were characterized. Results for individual mice are provided in Extended Data Fig. 7. **b,** The heavy- and light-mutations identified in panel (a) and characterized in Extended Data Fig. 8a were divided into those that improved neutralization for both isolates (red), those that improved it for only one indicated isolate (yellow or blue), and those that did not improve neutralization for either isolate (grey). The number of mutations characterized is indicated within each figure. **c,** Heavy-chain mutations of the four variants characterized in subsequent panels are listed with their aminoacid substitutions. **d,** A summary of IC_50_ values for wild-type 10-1074 and each 10-1074 variant is presented. Each point represents a mean of two independent sets of triplicates. Geometric mean values are indicated by a horizontal bar. Geometric mean values for 10-1074-sensitive isolates, for all isolates, and the breadth of each variant are indicated beneath the figure. Asterisks indicate significant differences between the indicated variant and wild-type 10-1074 (***p = 0.0008; **p = 0.0015; **p = 0.0027; repeated ANOVA followed by Dunnett’s multiple comparison tests). **e,** The polyreactivity of wild-type 10-1074 and the indicated 10-1074 variants including the rationally engineered 10-1074_GM_ were compared to polyreactive antibodies PGT128 and NIH45-46^W^ by immunofluorescence assays (IFA), using HEp-2 cells and 100 µg/ml of each antibody. Error bars indicate SD of mean intensity of three independent measurements. 10-1074 variants generated in this study (-y3, -v1, - v15) retained the low polyreactivity of wild-type 10-1074 (^ns^p = 0.8521; ^ns^p = 0.5496, ^ns^p = 0.7597; one-way ANOVA with Dunnett’s comparison tests). In contrast, 10-1074_GM_ was significantly more polyreactive than 10-1074 (***p = 0.0003). PGT128, and NIH45-46^W^ serve as positive controls, with significantly greater polyreactivity than 10-1074 (****p < 0.0001; ****p < 0.0001). Line indicates the value of a negative control sample provided by the manufacturer. Representative images of HEp-2 bound to the indicated antibodies are provided in Extended Data Fig. 8g. **f,** 5 mg/kg of the indicated 10-1074 variant was infused into 4-5 homozygous human FcRn Tg32 mice per group. Blood was drawn and antibody concentration was determined at the indicated time points. Error bars indicate SD. Asterisks adjacent to each point indicate significant differences between wild-type 10-1074 and the indicated variant, determined by mixed effects model of repeated measures followed by Dunnett’s multiple comparison tests (from earlier to later for each variant: 10-1074-y3: *p = 0.0184; 10-1074-v1: *p = 0.0251; *p = 0.0388; *p = 0.0236; *p = 0.0323; 10-1074-v15: *p = 0.0432; **p = 0.0051; 10-1074_GM_: *p = 0.0315; ***p = 0.0007; ***p = 0.0002; **p = 0.0083; **p = 0.0053). A second study with similar results is presented in Extended Data Fig. 8h.

Four of these 10-1074 heavy-chain mutations (N31K, E55D, V100dM, S100fA) and one light-chain mutation (M90L) potently enhanced neutralization of both CRF250 and BG505-T332N isolates (**Extended Data**). These five mutations were then combined in various ways and characterized their ability to neutralize BG505-T332N and YU2, the isolate originally used to select 10-1074^25^. We observed that 10-1074 variants with both V100dM and S100fA consistently neutralized these isolates more efficiently than wild-type 10-1074 (**Extended Data Fig. 8b**). V100d directly contacts a proteinaceous region in the third variable loop of Env as well as the glycan at asparagine 332. S100f also directly contacts this glycan (**Extended Data Fig. 8c**). We therefore characterized four additional 10-1074 variants, denoted 10-1074-y1 through-y4, each including these V100dM and S100fA mutations, against a wider isolate panel (**Extended Data Fig. 8d-e**). Each of these variants also neutralized every 10-1074-sensitive member of a global panel of HIV-1 isolates more efficiently than 10-1074 itself, with significantly lower geometric mean IC_50_ values. Among these variants, 10-1074-y3 was modestly more potent. We then compared 10-1074-y3 with 10-1074-v15, the most potent naturally emerging variant characterized in Fig. 5 (**Fig. 6c-d; Extended Data Fig. 8f**). We also included in these studies 10-1074-v1, which shares two residues with 10-1074-v15, one of which encodes novel glycosylation motif. All three variants neutralized a global panel of isolates significantly more efficiently than wild-type 10-1074 (**Fig. 6d**). Among these, 10-1074-y3 was significantly more potent than the other variants.

### Naturally emerging variants maintain the bioavailability of wild-type 10-1074

A key challenge with *in vitro* evolution and structure-based improvements of antibodies is the frequent loss of bioavailability associated with potency-enhancing mutations^30–33^. We hypothesized there is ongoing counter-selection *in vivo* against BCRs that react with self-proteins, or those which are more prone to proteolysis or aggregation. We therefore compared the polyreactivity and half-life of wild-type 10-1074, 10-1074-y3, the two naturally emerging 10-1074 variants (v1, v15), and the previously described rationally designed 10-1074 variant, 10-1074_GM_^25^. The polyreactivity of all three 10-1074 variants was similar to or lower than wild-type 10-1074, and significantly lower than 10-1074_GM_ and two polyreactive bNAbs (NIH45-46^W^, PGT128)^34–36^ (**Fig. 6e; Extended Data Fig. 8g**). 10-1074-y3 and both naturally emerging variants exhibited half-lives similar to or greater than the already long half-life of unmodified 10-1074^37,38^ in human FcRn transgenic mice, and markedly greater than 10-1074_GM_ (**Fig. 6f; Extended data Fig. 8h**). Notably, both naturally emerging variants exhibited the longest half-lives across two independent studies. We conclude that 10-1074 variants emerging from *in vivo* selection retained the low polyreactivity and improved the long half-life of 10-1074. These data suggest that *in vivo* evolution of native-loci edited B cells can select for antibodies with higher affinity without impairing their bioavailability.

### Affinity maturation of anti-SARS-CoV-2 antibodies *in vivo*

To extend this approach to additional antibodies, we used native-loci editing to generate B cells expressing two well-characterized SARS-CoV-2 neutralizing antibodies, ZCB11^39^ and S309^40^, which recognize the spike protein receptor-binding domain (RBD). Both antibodies are less potent against recent Omicron strains than earlier strains, and ZCB11 is fully resistant to BQ and XBB Omicron lineages^41,42^ (see **Extended Data Fig. 9a for RBD differences among these strains**). Roughly 4–6% primary B cells were edited to express each antibody, as indicated by their binding to the ancestral RBD (**Fig. 7a**). These edited cells were then characterized for their ability to bind monomeric RBD forms of two recent Omicron variants, BA.5 and XBB.1.5. ZCB11 bound only the BA.5 RBD, with lower affinity than to the ancestral RBD. The broadly neutralizing antibody S309, originally isolated from a SARS-CoV-1-infected person^40^, bound both BA.5 and XBB.1.5 RBDs, again with lower affinity than the ancestral SARS-CoV-2 RBD. To improve their neutralization potencies against these Omicron lineages, we engrafted mice with B cells engineered to express either ZCB11 or S309 and immunized them with antigens based on BA.5 or XBB.1.5, respectively. Specifically, engrafted mice were immunized twice with mRNA lipid nanoparticles expressing these membrane-associated RBDs modified with glycans to increase stability^43^ (**Extended Data Fig. 9b**). In contrast to HIV-1 Env immunogens (**Fig. 2b-d**), mRNA-expressed RBD vaccines elicited potent neutralizing plasma in unengrafted mice, and thus differences in plasma potency between engrafted mice and these control mice were less pronounced (**Fig. 7b; Extended Data Fig. 9c**). Nonetheless, CD45.1 donor cells expressing each antibody underwent clear hypermutation on both heavy and light chains (**Fig. 7c, Extended Fig. 9d**). We then characterized the most common ZCB11 and S309 variants that included substitutions or indels present in more than one mouse (**Fig 7d; Extended Data Fig. 9e**). These antibody variants bound proteins derived from their respective immunogen strains – BA.5 S protein for ZCB11 and XBB.1.5 RBD for S309 – with significantly greater affinity than unmodified forms of these antibodies, indicating that they are products of *in vivo* affinity maturation. To determine if this affinity maturation resulted in improved neutralization, we focused on the ZCB11 variant that bound the BA.5 S protein with the highest affinity (**Fig. 7d-e**). This variant, ZCB11-v9, bears a single L47P light-chain substitution (**Extended Data Fig. 9e**). ZCB11-v9 neutralized its immunogen strain BA.5 with 60-fold greater potency than unmodified ZCB11, and markedly enhanced the neutralization against the BA.5 Omicron precursor BA.2, as well as ancestral and several other pre-Omicron variants (**Fig. 7f-g**). It also neutralized the fully ZCB11-resistant BQ.1.1 Omicron strain, demonstrating that *in vivo* affinity maturation can result in increased antibody breadth. Interestingly, ZCB11 and S309 light chains both derive from the same variable segment, Vk3-20, and light-chain L47P was the most prominent light-chain mutation in one S309-engrafted mouse (**Extended Data Fig. 9d**). We accordingly characterized an S309 variant, S309-v13, whose light chain has been modified to L47P. This mutation improved S309 neutralization of the immunogen strain XBB.1.5, as well as XBB.1.16, and the immediate progenitor to both, XBB.1 (**Fig. 7g; Extended Data Fig. 9f**). S309-v13 was also more potent than S309 against the SARS-CoV-1 S protein and several pre-Omicron SARS-CoV-2 strains, but the L47P mutation impaired neutralization of three additional Omicron variants. The mechanism by which L47P improves the potency of both ZCB11 and S309 against multiple variants is unclear, but its location suggests that it might stabilize the interface between the heavy- and light-chain or subtly modify the conformation of the heavy-chain CDR3^39,40^. In summary, edited B cells expressing ZCB11 and S309 antibodies affinity matured *in vivo*, resulting in more potent and broader antibodies variants.

**Fig. 7:**
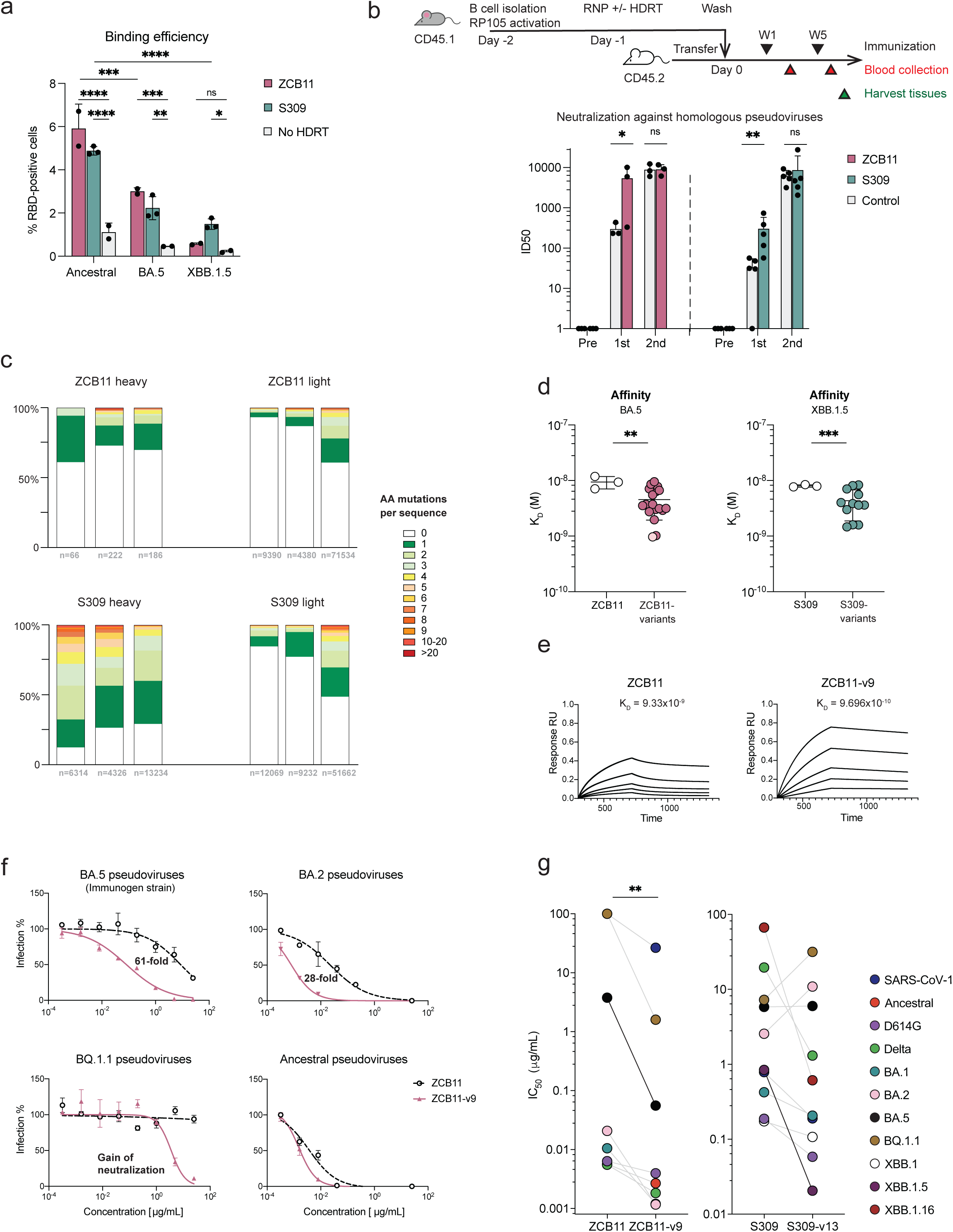
Affinity maturation of SARS-CoV-2 neutralizing antibodies. **a,** Primary murine B cells were native-loci edited as represented in Fig. 1a so that the heavy and light chains of the SARS-CoV-2 neutralizing antibodies ZCB11 or S309 were introduced into their corresponding murine loci. Edited cells were analyzed by flow cytometry using the S-protein receptor-binding domain (RBD) of the indicated SARS-CoV-2 variants conjugated to APC. Control cells were electroporated with the same RNP complex, without HDRT (no HDRT). Numbers indicate the percentage of RBD-positive cells. Each bar indicates the average of two to three independent experiments, and error bars represent SD. Significance was determined by two-way ANOVA followed by Tukey’s multiple comparison tests, from left to right: ****p < 0.0001; ****p < 0.0001; ***p = 0.0005; ***p = 0.0002; **p = 0.0022; ****p < 0.0001; ^ns^p = 0.7093; *p = 0.0197. **b,** Top panel presents schedule of immunization and blood collection used to generate plasma and cells analyzed in panels (c-d). Murine B cells were obtained from C57BL/6 mice marked with the CD45.1 allotype, engineered by native-loci editing, and adoptively transferred to CD45.2-positive recipient mice. These recipient mice were immunized with lipid nanoparticle (LNP)-encapsulated RNA expressing glycan-modified RBD fused to a transmembrane domain (gRBD-TM) at weeks 1 and 5. Mice engrafted with B cells expressing ZCB11 were immunized with mRNA-expressing the BA.5 gRBD-TM. Those engrafted with S309-edited B cell were immunized with XBB.1.5 gRBD-TM mRNA-LNP. Blood was collected at weeks 3 and 6. Bottom panel presents plasma neutralization of pseudoviruses of the respective immunogen strains. ‘Pre’ indicates a pool of plasma from mice before engraftment and immunization. Control indicates identically vaccinated mice without adoptive transfer. Error bars represent SD from three (ZCB11) or five (S309) mice. Significance was determined by mixed effects model of repeat measures followed by Šídák’s multiple comparison tests, from left to right: *p = 0.0320; ^ns^p > 0.9999; **p = 0.0016, ^ns^p > 0.9999. **c,** CD45.1 donor B cells from spleen and lymph nodes in ZCB11 or S309 recipient mice characterized in panel (b) were sequenced one week after their final vaccination. The number of amino-acid mutations per unique sequence is represented with proportions shown for each mouse. Each antibody and locus are indicated above the figure, and the number of unique sequences per mouse is shown below. **d,** Heavy and light chain variants with amino-acid substitutions found in multiple mice were expressed and characterized by bilayer interferometry (BLI) for their ability to bind the RBDs of their respective immunogens. Specifically 17 ZCB11 variants were analyzed for their ability to bind the BA.5 S protein and 12 S309 variants were analyzes using the XBB.1.5 RBD. The highest affinity ZCB11 variant, ZCB11-v9, studied in subsequent panels, is indicated by a lighter red. The affinity of these variants was compared to the average of three independent measurements using the respective wild-type forms of these antibodies. Error bars represent SD. Significance was determined by unpaired two-tailed t tests with Welch’s correction: **p = 0.0047; ***p = 0.0003. **e,** BLI binding curves representative of those used to generate panel (d), comparing unmodified ZCB11 and ZCB11-v9. K_D_ values are shown above each panel. **f,** Representative curves comparing ZCB11 (grey) and ZCB11-v9 (red) neutralization against selected SARS-CoV-2 isolates, including the BA.5 immunogen strain, are presented. Fold improvement is indicated each panel. Error bars indicate SEM from triplicates. **g,** Each point represents the IC_50_ values for a panel of SARS-CoV-1 or −2 variants of wild-type ZCB11 and S309 as well as ZCB11-v9 and S309-v13, generated as in panel (f) and Extended Data Fig. 9f. Lines connect the same SARS-CoV-1 or −2 variant, and dark line connects the immunogen strain used to generate antibody variants (BA.5 for ZCB11, XBB.1.5 for S309). Each point represents a mean IC_50_ of two independent sets of triplicates. Significance was determined by paired two tailed t tests: **p = 0.0047.

## Discussion

Here we present an efficient method for reprogramming murine B cells to express exogenous B cell receptors. We call this reprogramming technique ‘native-loci editing’ because the recombined murine heavy- and kappa-chain variable regions are simply replaced by their human counterparts, without displacement or additional regulatory elements. We achieve this by simultaneously introducing a double-strand break at the 3’-most J segment of each locus. We then template repair with homology arms complementary to variable-segment promoters located near the 5’ of these loci and to introns immediately downstream of these J-region breaks. In successful cases, the resulting BCRs are expressed from endogenous promoters, and edited cells are indistinguishable from the results of a VDJ (or VJ) recombination event, except that an exogenous human variable gene replaces the endogenous murine gene. One notable aspect of this approach is that a single double-stranded break at each locus facilitated excision of long genomic regions, often greater than 100 kb, as previously reported for induced pluripotent stem cells^44^. The spatial arrangement of the BCR loci, which promotes long-range interactions necessary for VDJ(VJ) recombination^45^, may have contributed to the efficiency of these long excisions and their replacement with human variable genes.

Earlier pioneering studies^12–16^ employed an alternative method in which a cassette – expressing an exogenous promoter, light-chain variable and constant gene segments, heavy-chain variable segments, and several other regulatory sequences – was introduced into an intron downstream of the 3’-most mouse JH gene. We compared our native-loci method to one such intron-targeting approach^12^ by introducing the same number of B cells engineered through each technique into wild-type mice, and then vaccinating these mice with appropriate antigens. We observed higher neutralization activity in plasma of mice engrafted with native loci-edited B cells. We also observed significantly higher levels of class switching and SHM in these cells. Critically, efficient SHM facilitated affinity maturation in B cells edited to express the HIV-1 bNAb 10-1074 and the anti-SARS-CoV-2 antibodies ZCB11 and S309. Robust SHM and affinity maturation of edited BCRs suggest that native-loci editing does not significantly disrupt development or function of the edited B cell. In contrast, we observed lower SHM and diversity in mice engrafted with B cells edited using an intron-targeting approach. These lower SHM frequencies are consistent with previous reports^14,15,17^ and suggest that elements of the intron-targeting cassette interfere with processes necessary for a potent and diverse antibody response. We note a similar observation with chimeric-antigen receptor (CAR) T cells. Specifically, CARs expressed from an endogenous T cell receptor alpha-chain promoter controlled tumor progression *in vivo* more efficiently than those expressed from an exogenous promoter^46^. Both results suggest that minimal perturbation of the endogenous regulatory apparatus results in greater preservation of lymphocyte function.

Because a native-editing approach better preserves B-cell activities *in vivo*, it expands the usefulness of B-cell engineering for at least four applications. First, engineered B cells can transform wild-type mice into a useful tool for evaluating vaccines. This application may be especially useful in the case of HIV-1, where the most promising strategies attempt to affinity mature bNAbs from their human precursors. We show here that mice engrafted with B cells expressing bNAbs against two key Env epitopes raise potent neutralizing plasma. It would therefore be straightforward to engraft cells expressing precursors to these antibodies and monitor their response to candidate antigens and vaccine strategies. Native-loci editing generates more robust B cell responses than other editing approaches, enabling more precise comparisons of these strategies. Unlike knock-in mice now used for this purpose, engineering of naïve mature B-cells requires a shorter development cycle, enables introduction of antibody combinations and libraries of variants, and can be implemented in many species.

Second, native-loci editing can also be used to address open scientific questions in B-cell biology. For example, by introducing defined input sequences, it can provide insight into the processes and patterns of somatic hypermutation. It can also help delineate principles of peripheral tolerance and determine whether expression of a BCR from a naïve mature cell is itself tolerizing.

Third, robust affinity maturation also expands the potential therapeutic applications of engineered B cells if safety concerns can be addressed. Engineered B cells can provide high levels of therapeutic antibodies for much longer than passive infusion, and they are likely to elicit fewer anti-drug antibody responses than with gene-therapy delivery systems^47,48^. The ability of native-loci edited cells to affinity mature is especially useful in the case of an established HIV-1 infection, enabling them to adapt to the specific proviruses reactivated in HIV-1-positive persons undergoing treatment interruption. To fully control an established infection, multiple Env epitopes must be targeted to ensure that suppression is consistent and lasting. We show here that B cells edited to express 10-1074 can expand and hypermutate alongside those edited to express VRC26.25-y. Both sets of cells responded to the same Env antigen independently, suggesting that key Env epitopes can be recognized simultaneously with adaptive BCRs. This same approach might be applied in special cases to other rapidly evolving pathogens, for example SARS-CoV-2, with cells expressing broad but hard-to-elicit neutralizing antibodies.

Finally, as we demonstrated here, native-loci editing of B cells provides an alternative to both *in vitro* display approaches and transgenic mice for improving therapeutic antibodies. We observed that nearly every 10-1074 variant directly isolated from engrafted mice bound the homologous antigen with higher affinity and neutralization potency than wild-type 10-1074. We further observed that the majority of mutations found in multiple mice improve the potency of 10-1074, and that these mutations could be combined to generate a variant with more than twice the potency against a global panel of isolates. Notably, none of these individual mutations or the selected heavy chains significantly impaired neutralization, implying that *in vivo* selection effectively eliminates less potent variants. Similarly, we showed that two SARS-CoV-2 neutralizing antibodies affinity matured in response to RBD antigens they initially bound weakly. Most clearly, the class II antibody ZCB11 matured to neutralize the BA.5 Omicron strain with 60-fold greater efficiency, and in the process gained the ability to neutralize the previously resistant BQ.1.1 strain.

Therefore *in vivo* affinity maturation can complement more conventional *in vitro* methods for improving antibody affinity like phage-, yeast-, and mammalian cell-display techniques^49–51^. Each has different strengths. These *in vitro* methods often start with much larger libraries, but the continuous diversification and coordinated selection process in germinal centers may more effectively explore useful sequence space over time. Also, diversified HDRT libraries can further improve sampling of sequence space *in vivo*, combining the strengths of i*n vitro* and *in vivo* evolution. More importantly, antibodies that emerge from *in vivo* selection may have additional properties useful to human therapeutics. For example, mammalian immune systems can select against BCRs that express inefficiently, aggregate, recognize self, or are easily proteolyzed *in vivo*. Thus, the naturally emerging 10-1074 variant characterized in pharmacokinetic studies exhibited an *in vivo* half-life comparable to or greater than 10-1074, which already has one of the longest half-lives of HIV-1 bNAbs. In contrast, a previously described rationally designed 10-1074_GM_ exhibited higher polyreactivity and shorter half-life, one of many examples whereby potency improvements generated through *in vitro* technique*s* impaired bioavailability^30–33^. In short, *in vivo* affinity maturation uniquely selects against antibodies with properties that would limit their therapeutic efficacy. Another *in vivo* approach for improving antibodies utilizes transgenic mice, often to express a single bNAb or bNAb precursor. As mentioned, compared with these knock-in mice, B-cell engineering can be implemented more rapidly, accommodates variant libraries, and can be used in other species. In addition, unlike knock-in mice, the engineered B cells bypass central tolerance, perhaps enabling expression of antibodies that would be clonally deleted in transgenic mouse models^52–55^.

Despite the promise of native loci-editing technology, this study has several limitations. First, although *in vivo* affinity maturation increased 10-1074 affinity for a homologous SOSIP by 10-fold, the effect on HIV-1 neutralization using a heterologous panel of global isolates was more modest, and our most potent variant combined mutations that were not observed together *in vivo*. These relatively small potency improvements may reflect the high initial potency of 10-1074 against the homologous BG505-T332N envelope glycoprotein, perhaps also explaining why our limited efforts to identify improved variants of the highly potent VRC26.25-y did not yield significant results. However, we observed more robust improvement with the anti-SARS-CoV-2 antibody ZCB11 perhaps because its initial affinity with the Omicron BA.5 RBD antigen was lower. Further improvements might be obtained with a more diverse set of immunogens, more effective vaccines, greater initial BCR diversity, and appropriate coupling of heavy and light chains. Second, we are using mice to counter-select against properties that impair bioavailability but some of these properties – for example interactions with specific self-proteins – may differ between mice and humans. Extension of this approach to a second animal model, including non-human primates, will mitigate this concern. Third, comparisons between native-loci and intron editing were made using one intron-based approach, and our specific implementation of this approach may not be fully optimal. Nonetheless, to date, no examples of robust or useful affinity maturation have been reported with this method, implying that intron-editing impairs B cell function. Finally, although we have demonstrated greater functionality of native-edited B cells over those generated with an intron-based approach, *ex vivo* culture, electroporation, and Cas12a-mediated editing may nonetheless perturb these cells in subtle ways. Moreover, as with most CRISPR-engineered cells, off-target activities carry the risk of cell transformation. These concerns are especially relevant to potential therapeutic applications of engineered B cells. Further study of the safety and functionality of these cells, and further improvements in editing technologies, will therefore be necessary before they can be used in humans. The potential of these cells to adaptively control chronic infections should help motivate these efforts.

In summary, we have developed methods to directly replace recombined heavy- and kappa-variable genes with those from human antibodies, leaving each locus otherwise unmodified. Compared with earlier approaches, this native-loci editing approach resulted in more potent neutralizing plasma, more efficient SHM, and importantly, robust *in vivo* affinity maturation. This affinity maturation guided the development of more potent 10-1074 variants without loss of bioavailability, and generated broader, more potent variants of two SARS-CoV-2 antibodies. Because this approach better preserves the *in vivo* activities of engineered B cells, it can be used to improve the properties of therapeutic antibodies, evaluate conventional vaccines, address previously inaccessible questions in B-cell biology, and perhaps develop new B cell-based therapies.

## Acknowledgements

The authors would like to gratefully acknowledge Dr. Hyeryun Choe for her advice and careful reading of the manuscript. Funding is provided by National Institutes of Health awards U19 AI149646 (M.F.); R21 AI152836 (M.F.); R01 DA056771 (M.F.).

## Author contributions

Y.Y and M.F. conceived of the study; Y.Y. designed, performed, and analyzed experiments; H.P. assisted with surface plasmon resonance studies; Y.Y., Y.G., B.Q., L.Z., W.H. and T.O. developed key reagents or provided useful information; Y.J. and C.C.B. provided computational analysis and advice; Y.Y. and G.C. performed statistical analyses; M.F. provided funding support; Y.Y. and M.F. wrote the manuscript.

## Supplementary information

Methods

Extended Data Fig. 1 to 9

Supplementary table 1

